# Cross-sector collaboration is more effective than single sector actions at mitigating SARS-CoV-2 in white-tailed deer

**DOI:** 10.1101/2023.10.13.562192

**Authors:** Jonathan D. Cook, Elias Rosenblatt, Graziella Direnzo, Evan H. Campbell Grant, Brittany A. Mosher, Fernando Arce, Sonja Christensen, Ria R. Ghai, Michael C. Runge

## Abstract

One Health helps achieve optimal health outcomes for people, animals, plants, and their shared environments. We describe a multidisciplinary effort to better understand and mitigate SARS-CoV-2 spread in white-tailed deer across One Health sectors. We first framed the risk problem with three governance sectors that manage captive and wild deer and human public health. The framing included the objectives for each sector, interactions that facilitate human-to-deer and deer-to-deer transmission, and alternatives intended to reduce risk. We then developed a dynamic compartmental model that linked wild and captive deer herds and humans and simulated SARS-CoV-2 dynamics. For baseline conditions, we estimated that median SARS-CoV-2 prevalence in wild and captive herds varied between 0.03 – 0.07, incidence between 0.68 – 1.46, and probability of persistence between 0.64 – 0.97 across 120-day simulations. We then tested single-sector alternatives alone and in combination with other sector actions. We found that single sector alternatives varied in their ability to reduce transmission and that the best performing alternative required collaborative actions among wildlife management, agricultural management, and public health agencies.

## 1. Introduction

The emergence and global spread of severe acute respiratory syndrome coronavirus 2 (SARS-CoV-2) in humans, domestic animals, and wildlife has created a challenging One Health disease problem. The virus first evolved in a mammalian host, likely Old-World bats of the family *Rhinolophidae*^1^, but its transmission to and subsequent spread among the human population has propelled a pandemic of considerable economic, social, and cultural significance. Widespread human transmission has resulted in spillback to numerous mammalian hosts including domestic, farmed, and wild animals (U.S. Department of Agriculture SARS-CoV-2 Dashboard accessed 30-Jun-2023^2^). Infection of farmed mink (*Neogale vison*) and white-tailed deer (deer, *Odocoileus virginianus)* have caused particular concern among the public health community because of the documented animal-human transmission and emergence of novel variants.

Detections of SARS-CoV-2 in deer across four U.S. states began in 2020^3^. Additional surveillance has repeatedly detected multiple strains that are closely related to dominant strains circulating in nearby human populations at the same times^4,5,6^. In 2022, a highly divergent lineage with mutations associated with non-human hosts was detected in deer and a person with known contact with deer in Ontario, Canada^7^. Legacy strains have also been detected in deer months after they were largely supplanted by new variants within the human population^7^. These findings indicate that human-to-deer transmission has likely occurred repeatedly, and that deer can transmit and sustain SARS-CoV-2 among their populations^6^. Taken together, it is possible that deer are SARS-CoV-2 reservoirs, and that they may be a source of novel variants that threaten public health by evading diagnostic detection or affecting therapeutic and vaccine efficacy.

Like other zoonoses, managing SARS-CoV-2 transmission between human and non-human hosts is a challenge that requires a One Health approach. Deer are widespread and abundant in diverse settings across North America, resulting in many possible exposure routes between humans and deer. Furthermore, the U.S. management agencies’ governance structure is complex with disjunct regulatory authorities that could challenge effective disease management. Understanding linkages across One Health sectors (human, wildlife, and agriculture), and identifying whether coordinated responses are necessary for mitigating disease transmission is a considerable challenge.

An evidence-based approach to inform risk management decisions is essential for effective mitigation. Therefore, we first coordinated a multi-jurisdiction team of decision makers in the U.S. with authority to manage wild deer, captive deer, or human public health. This decision framing included articulation of sector-specific and shared fundamental objectives, identification of causal chains of interactions that may facilitate spillover and spread, and a specification of possible management alternatives. Second, we developed and used a deterministic compartmental model to evaluate spillover and spread rates in isolated deer herd settings, and to estimate prevalence, incidence, and persistence in a range of connected human-deer scenarios. Here, we focus on evaluating the decision space of the One Health collaborative using a model that was developed and described in detail in Rosenblatt et al. (2023)^8^. Collectively, our decision analysis and risk modeling efforts can guide agencies with purview over risk mitigation to make data-driven, transparent, and reproducible decisions that help meet their fundamental objectives.

## 2. Methods

### 2.1 Overview

We sought to frame risk mitigation decisions using a decision analytical approach^9,10^. While its application to One Health issues is novel, framing multi-sector disease problems collaboratively improves clarity of decision-making within sectors, identifies opportunities for resource sharing across sectors, and may allow for creative alternatives that enhance One Health outcomes. We convened a guidance committee composed of three sectors involved with deer management and human public health (Supplementary Table 1). It included seven members from wildlife management agencies (WM), five from agricultural management agencies (AM), and five from human public health agencies (PH). We then used a series of 10 meetings to frame the decision context and identify a set of alternatives for evaluation.

### 2.2 Components of decision framing

The first step was to articulate the scope of the problem including the spatial extent and specific authorities of agency representatives. The guidance committee defined the spatial scope of the evaluation as regions in the U.S. with high wild deer density, where captive deer occur, and with human populations in proximity to both wild and captive deer (e.g., agro-forested midwestern states). In terms of authorities, state wildlife management agencies in the U.S. have primary authority to manage wild deer within their boundaries, however other federal agencies have shared authority on certain federal lands. In most U.S. states, captive deer are managed by state agricultural agencies, including farmed deer and deer on exhibition. In some locations these authorities are shared with state wildlife management agencies. Animals kept in captivity and those that are involved in trade are regulated in a manner that is consistent with their intended use. Public health agencies make decisions to promote health at all governance levels, including state, tribal, local, and territorial health departments. In general, public health agencies use their authorities to provide voluntary health recommendations and to communicate zoonotic disease detections so that at-risk groups can practice individual risk mitigation.

We then worked on framing the specific interests (i.e., objectives), the causal chains of interactions that may affect disease transmission, and interventions to interrupt transmission. We summarized this information using an influence diagram to map connections among fundamental objectives and alternatives following the causal chains^11^.

The guidance committee identified eight fundamental objectives to consider in this context. In general, the representatives from public health expressed interest in minimizing the morbidity and mortality of COVID-19 in humans by limiting infection. Wildlife and agricultural management agencies expressed interest in maintaining individual and population health of deer in support of the North American model of wildlife conservation^12^ and the Animal Welfare Act (7 U.S.C. 2131 et seq.). Relatedly, all agencies were interested in limiting SARS-CoV-2 transmission within deer herds and other hosts because of the potential for viral strain mutation, recombination with endemic coronaviruses, and the emergence of novel variants. Novel variants can change the host range, transmissibility, and other viral properties, potentially resulting in new waves of infection within human populations. Thus, efforts to limit disease transmission among deer, humans, and other wildlife species are important. In addition to the disease-related objectives, there were several other objectives expressed among the management sectors related to hunter satisfaction, business activities, and management authorities and public trust (Table 1).

**Table 1.** Fundamental objectives and measurable attributes for SARS-CoV-2 risk mitigation decisions in deer and across One Health sectors addressing COVID-19 in AM, PH, and WM. AM, Agricultural management agencies. PH, Public Health agencies. WM, Wildlife management agencies.

| Fundamental Objectives (Measurable Attribute) |
| --- |
| Fundamental Objective 1. Minimize SARS-CoV-2 infection in humans. Sectors: PH, AM.<br>(Measurable Attribute 1. Median SARS-CoV-2 incidence in deer across the 120-day period). |
| Fundamental Objective 2. Maximize the health of deer. Sectors: AM, WM.<br>(Measurable Attribute 2. SARS-CoV-2 incidence in deer across the 120-day period). |
| Fundamental Objective 3. Minimize risk of SARS-CoV-2 evolution. Sectors: PH.<br>(Measurable Attribute 3. Median probability of SARS-CoV-2 persistence in deer across the 120-day period) |
| Fundamental Objective 4. Maximize deer hunter satisfaction and participation. Sectors: AM, WM. |
| Fundamental Objective 5. Minimize harm to privately owned captive cervid businesses. Sectors: AM. |
| Fundamental Objective 6. Maintain the current management authority of wildlife management, agricultural, and public health agencies. Sectors: AM, PH, WM. |
| Fundamental Objective 7. Maximize public appreciation, trust, and acceptance of management. Sectors: AM, PH, WM. |
| Fundamental Objective 8. Provide timely and consistent data-driven measures of local zoonotic disease risks. Sectors: AM, PH, WM. |

The next step was to map the causal chain as an influence diagram. The guidance committee identified two distinct transmission modes for human-to-deer and deer-to-deer transmission: direct transmission (aerosol inhalation or fluids deposited during social contact), and indirect transmission (contact with contaminated fomites or wastewater) (Figure 1, central component). Within those two modes were six sources of introduction and spread in wild and captive deer (Figure 1, green rounded rectangles). For this evaluation, we focused on direct modes of human-to-deer and deer-to-deer transmission based on the highly social behaviors of deer that include nose-to-nose contact and experimental studies that indicate viral replication in the upper respiratory tract and shedding via nasal secretions^13^ (Figure 1, dotted line). There is currently no evidence to support indirect transmission modes.

**Figure 1.**
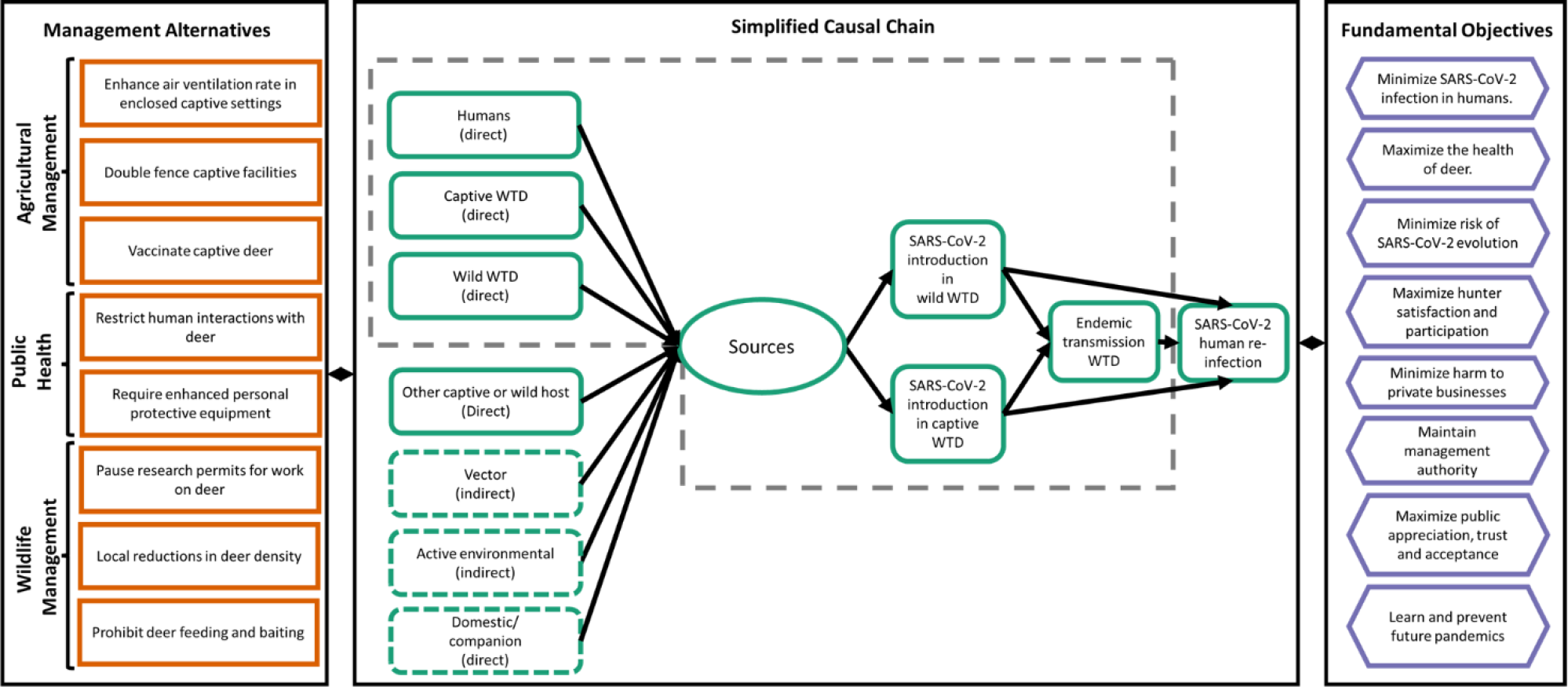
Influence diagram for SARS-CoV-2 introduction and spread in wild and captive deer. The green ovals show sources of SARS-CoV-2 introduction in wild and captive deer; the orange rectangles show the set of alternatives identified across sectors, and the purple hexagons show the collective set of fundamental objectives that each of the three One Health sectors may consider when making risk management decisions. The dotted line is the area of the causal chain that we focused on for this risk assessment.

Next, we used the diagram to generate a full set of alternatives (Supplementary Table 2), and then pared back the list to include a subset of alternatives of greatest interest of committee members. The selected alternatives for the AM sector included requirements to: 1) enhance air ventilation rate in enclosed agricultural/zoo/farm settings for captive deer, 2) double fence captive deer facilities, and 3) vaccinate captive deer (including booster dosing). Public health representatives considered two areas that could be the focus of education and training efforts: 1) encourage the public to limit interactions with wild deer in suburban settings (not related to hunting activities) like public parks, and private lands where deer may congregate (e.g., golf courses, backyards), and 2) encourage the proper use of enhanced personal protective equipment during interactions with captive and wild deer. For WM, the alternatives included: 1) a pause on permitted research involving close contact with wild deer, 2) reductions in wild deer density, and 3) prohibiting wild deer feeding and baiting.

We then developed a deterministic compartmental disease model that could test the efficacy of these alternatives in a simulated system where humans, wild and captive deer could interact (Rosenblatt et al. 2023 for details). The model allows captive and wild deer to transition among susceptible (s), infectious (i), and recovered (r) compartments before eventually returning to the susceptible compartment. The model estimates proportions of individuals in each disease compartment, does not include demography, and assume a closed population (i.e., no births, deaths, immigration, or emigration). Transmission occurs in deer from interacting with infectious humans or deer in captive and wild herds. Deer are assumed to be evenly distributed with all individuals subjected to the same contact rates. We parameterized the model to simulate the midwestern U.S. during the fall season (September-December; 120-day window) when humans may contact deer during hunting and other activities and acquired parameter estimates using empirical data and expert elicitation^8^ (Supplementary Table 3).

To explore variation in settings where humans and deer may interact, we evaluated wild deer in rural and suburban settings and captive deer in low- and high-density facilities (i.e., a total of four settings: rural, suburban, ranch, and intensive). For model simulation purposes, we defined a rural setting as a free-ranging deer population at a density of 10 deer/km^2[14]^ in a 26% forested landscape with human density of 3.1 humans/km^2^. For a suburban deer setting, we maintained the same deer density and forested cover, but we increased human density to 100 humans/km^2^ and human-deer contact rate. For the farmed deer setting, we defined a low-density condition equal to the density of rural wild deer but assumed an elevated human-deer contact rates (e.g., from supplemental feeding and other herd maintenance activities). We refer to this low-density captive deer setting as the ‘ranch’. Lastly, we defined a high-density setting (‘intensive’) as a herd in an enclosed environment and assumed less air exchange and elevated deer-to-deer and human-to-deer contact rates relative to the ranch setting (Supplementary Table 3).

To understand model performance and baseline conditions prior to evaluating alternatives, we first estimated disease introduction (from humans) and spread in the four settings. We ran 200 replicate simulations to sample over parametric uncertainty with no initial infections in deer and a continuous hazard of human-to-deer transmission. We summarized results as the probability of at least one human-to-deer SARS-CoV-2 transmission event in 120 days in a completely susceptible population of 1,000 deer and the basic reproduction number, *R*_0_ (i.e., average number of secondary cases from an average primary case in an entirely susceptible population^15^).

We then combined these settings into scenarios that allowed captive deer, wild deer, and humans to interact. We explored two scenarios: (1) interacting rural wild deer, farmed ranch deer, and rural human populations (rural x ranch), and (2) interacting suburban wild deer, captive intensive deer, and suburban human populations (suburban x intensive). Based on human-deer and deer-deer contact rates, we designed these combinations to explore a lower risk scenario (scenario 1) and a higher risk scenario (scenario 2). For an in-depth explanation of the model and its parameterization, see Rosenblatt et al. (2023)^8^ and Supplementary Methods for this manuscript.

To evaluate the performance of the alternatives on disease-related objectives (Objectives 1-3) in the two scenarios, we evaluated three different measurable attributes (Table 1). The measurable attribute for objective 1 was the median prevalence of SARS-CoV-2 in wild and captive deer. For objective 2, we used median incidence of SARS-CoV-2 in wild and captive deer. Finally, for objective three we used the median probability of persistence of SARS-CoV-2 in wild and captive deer. We defined persistence as any replicate with a proportion in the infectious compartment at equilibrium that exceeded 0.001. The goal was to identify alternatives that could minimize estimates of each measurable attribute. The complete set of results for each of the two scenarios are reported in a consequence table, which summarize the performance of each objective relative to the alternatives considered and help make tradeoffs visually apparent.

## 3. Results

We found that the average probability of at least one human-to-deer transmission event in a population of 1,000 deer and across the 120-day window was lower for wild deer than captive deer in each respective setting. The median probability of transmission was 0.01 (95% Prediction Interval [PI]: 2.2 × 10^-4^ – 0.25) and 0.20 (95% PI: 0.02 – 0.96) in rural and suburban wild settings, respectively (Figure 2). In captive settings, the probability of human-to-deer transmission was 0.55 (95% PI: 0.03 – 0.99) for ranch deer and 0.87 (95% PI: 0.08 – 1.0) for intensive captive deer (Figure 2). For deer-to-deer transmission within wild and captive herd settings, we found that captive herd settings were more likely to have sustained spread. The median estimate of *R*_*0*_ in intensive captive facilities was 6.24 (95% PI: 0.23 – 144.0) and in captive ranch facilities it was 1.53 (95% PI: 0.08 – 25.40, Figure 2). We found that the median *R*_*0*_ was less than 1.0 in both rural and suburban wild deer settings (median: 0.74 (95% PI: 0.05 – 13.3)); however, estimates of *R*_*0*_ were highly skewed with many parameter combinations resulting in values greater than 1.0 (Figure 2).

**Figure 2.**
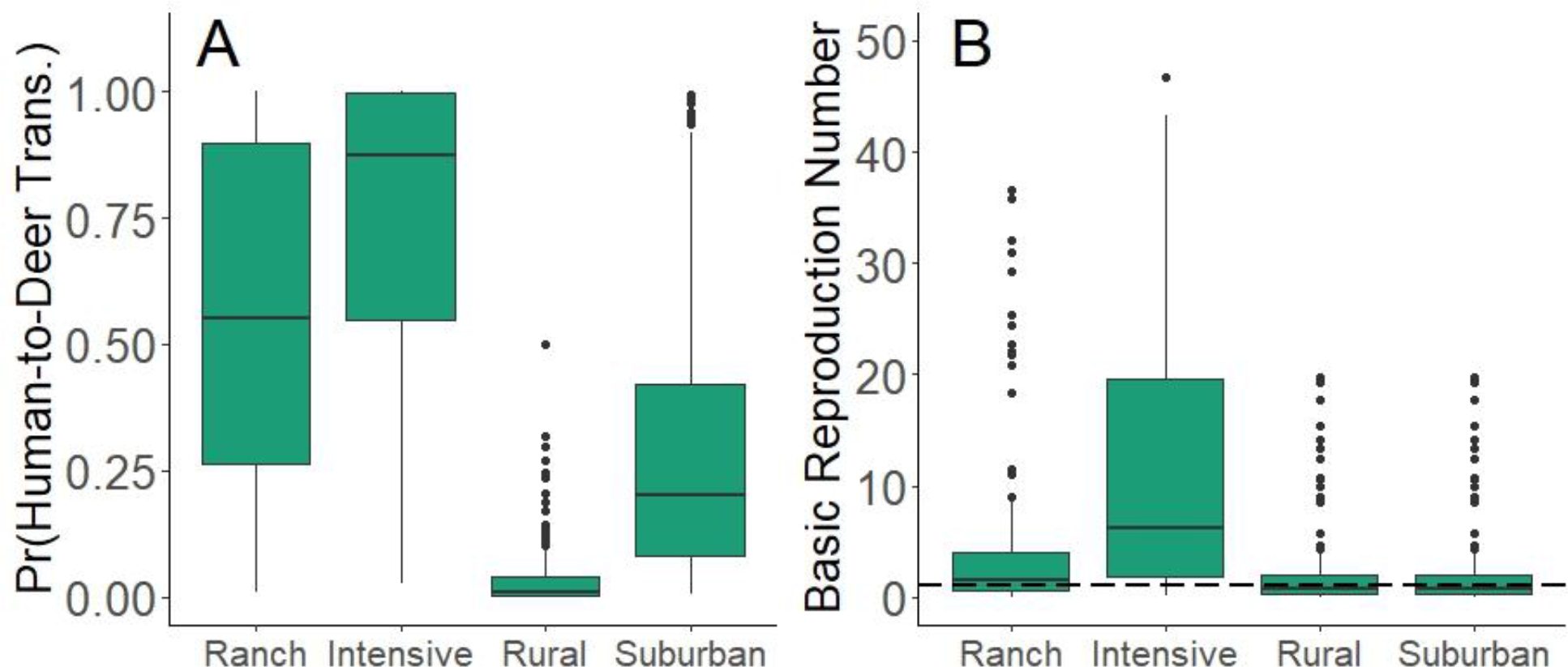
Probability of human-to-deer transmission and basic reproduction number for deer-to-deer transmission in four different captive and wild deer settings (Panels A - B): Boxplots of the probability of human-to-deer transmission (A) and basic reproduction number (B) in ranch and intensive captive deer herds and rural and suburban wild deer herds. Human-to-deer transmission and the basic reproduction number (*R*_*0*_) was typically higher in captive herd settings. We ran 200 replicate simulations to sample over parametric uncertainty with no initial infections in deer and a continuous hazard of human-to-deer transmission. Pr(Human-to-Deer transmission) is the probability of at least one human-to-deer SARS-CoV-2 transmission event in 120 days in a completely susceptible population of 1,000 deer and across the 200 replicate simulations. The basic reproduction number (B) is the average number of secondary cases from an average primary case in an entirely susceptible population.

For the scenarios, we found that baseline estimates of prevalence, incidence proportion (the average number of infections during the season), and probability of persistence varied between connected captive and wild deer herds according to setting types. In general, intensive captive herds interacting with suburban deer and humans had the highest overall prevalence (median: 7%; 95% PI: 0.00 – 17.90%), whereas rural wild deer interacting with captive ranch deer and humans had the lowest prevalence (median: 3%, 95% PI: 0.00 – 11.10%) (Figure 3; Table 2). For incidence proportion, intensive captive herds again had the highest estimate of 146% (i.e., the average deer was infected 1.46 times across the 120-day window; 95% PI: 0.01 – 372%)) and rural, wild deer had the lowest estimate (68%; Figure 3; Table 2; 95% PI: 0.00 – 231%). For the probability of persistence, intensive captive facilities were most likely to sustain spread of SARS-CoV-2 (97% of simulations) whereas wild, rural deer were the least likely (64% of simulations; Table 2).

**Figure 3.**
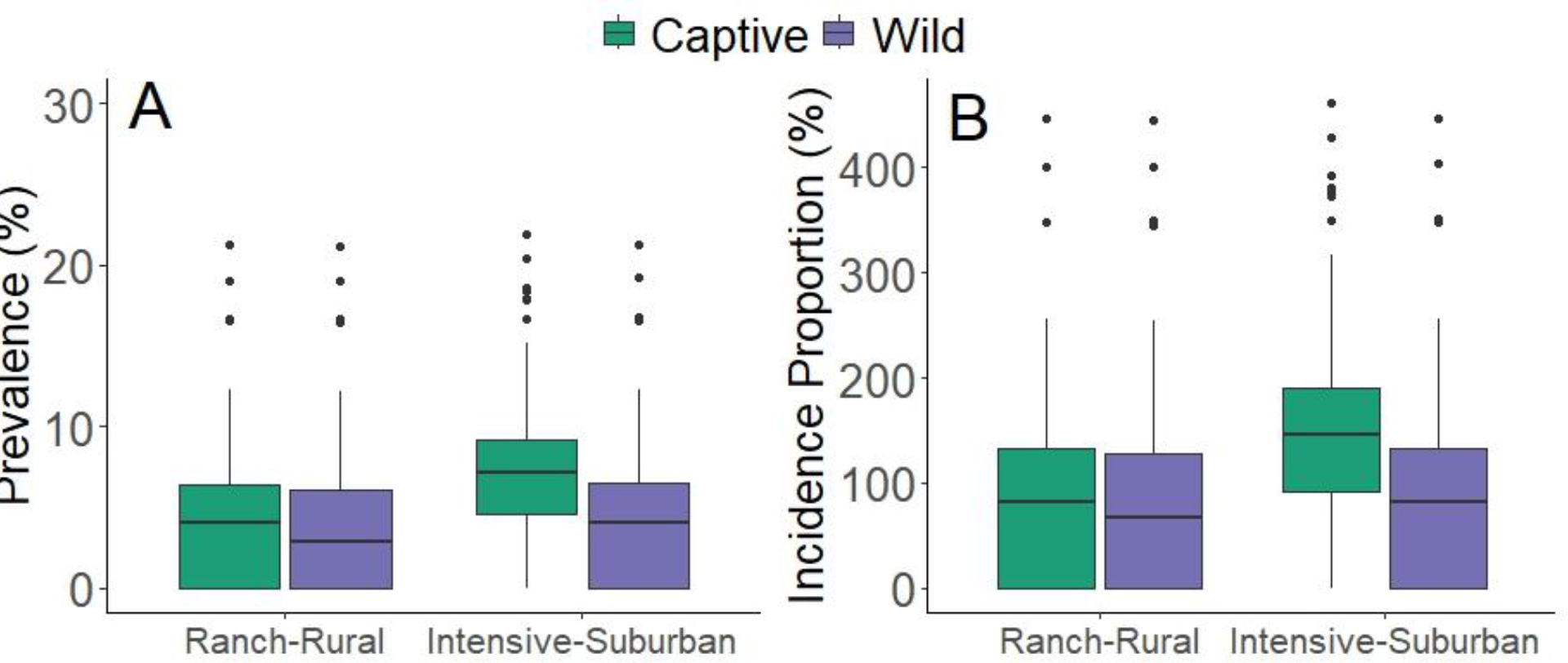
Deer-to-deer SARS-CoV-2 prevalence and incidence proportion in the two scenarios (Panels A - B): Boxplots of prevalence (A) and incidence proportion (B) in two scenarios of interacting wild and captive deer herds. prevalence in deer was used as a measure of human health risk, which related to Fundamental Objective 1, and incidence proportion was used a as a measure of deer health risk, which related to Fundamental Objective 2.

**Table 2.**
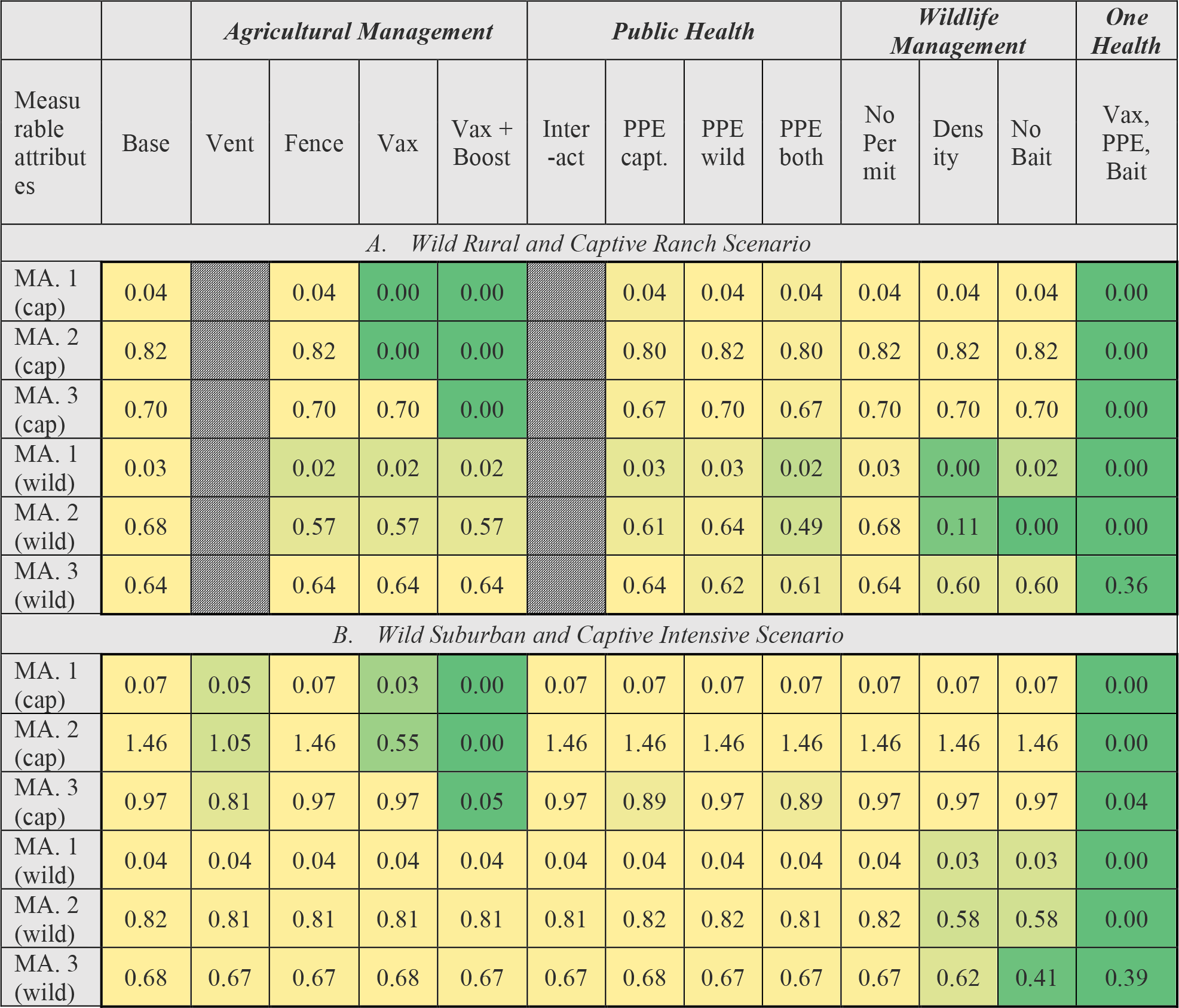
Consequence table showing the effectiveness of sector specific and One Health alternatives on achieving the fundamental objectives for the two scenarios. Green shades indicate better performing alternatives for each of the six measurable attributes. Yellow shades indicate worse performing alternatives and color gradients indicate relative performance of alternatives across each row (attributes). The One Health alternatives (i.e., joint actions taken by all sectors simultaneously) led to the best outcomes. Crosshatched cells indicate those where we did not evaluate those alternatives. Abbreviations: Base=Baseline, Cap=Captive, Density= Reduce deer density, Fence= Double fence captive deer facilities, MA1= median prevalence of SARS-CoV-2, MA2= median incidence of SARS-CoV-2, MA3= median probability of persistence of SARS-CoV-2, No Bait=Prohibit feeding and baiting, No Permit= Pause of permitted research involving close contact with wild deer, Interact=Reduced human-wild deer interactions, Vent=Enhance air ventilation rate, Vax= Vaccinate captive deer

|  |  | *Agricultural Management* |  |  |  | *Public Health* |  |  |  | *Wildlife Management* |  |  | *One Health* |
| --- | --- | --- | --- | --- | --- | --- | --- | --- | --- | --- | --- | --- | --- |
| Measurable attributes | Base | Vent | Fence | Vax | Vax + Boost | Inter-act | PPE capt. | PPE wild | PPE both | No Permit | Density | No Bait | Vax, PPE, Bait |
| *A. Wild Rural and Captive Ranch Scenario* |  |  |  |  |  |  |  |  |  |  |  |  |  |
| MA. 1 (cap) | 0.04 |  | 0.04 | 0.00 | 0.00 |  | 0.04 | 0.04 | 0.04 | 0.04 | 0.04 | 0.04 | 0.00 |
| MA. 2 (cap) | 0.82 |  | 0.82 | 0.00 | 0.00 |  | 0.80 | 0.82 | 0.80 | 0.82 | 0.82 | 0.82 | 0.00 |
| MA. 3 (cap) | 0.70 |  | 0.70 | 0.70 | 0.00 |  | 0.67 | 0.70 | 0.67 | 0.70 | 0.70 | 0.70 | 0.00 |
| MA. 1 (wild) | 0.03 |  | 0.02 | 0.02 | 0.02 |  | 0.03 | 0.03 | 0.02 | 0.03 | 0.00 | 0.02 | 0.00 |
| MA. 2 (wild) | 0.68 |  | 0.57 | 0.57 | 0.57 |  | 0.61 | 0.64 | 0.49 | 0.68 | 0.11 | 0.00 | 0.00 |
| MA. 3 (wild) | 0.64 |  | 0.64 | 0.64 | 0.64 |  | 0.64 | 0.62 | 0.61 | 0.64 | 0.60 | 0.60 | 0.36 |
| *B. Wild Suburban and Captive Intensive Scenario* |  |  |  |  |  |  |  |  |  |  |  |  |  |
| MA. 1 (cap) | 0.07 | 0.05 | 0.07 | 0.03 | 0.00 | 0.07 | 0.07 | 0.07 | 0.07 | 0.07 | 0.07 | 0.07 | 0.00 |
| MA. 2 (cap) | 1.46 | 1.05 | 1.46 | 0.55 | 0.00 | 1.46 | 1.46 | 1.46 | 1.46 | 1.46 | 1.46 | 1.46 | 0.00 |
| MA. 3 (cap) | 0.97 | 0.81 | 0.97 | 0.97 | 0.05 | 0.97 | 0.89 | 0.97 | 0.89 | 0.97 | 0.97 | 0.97 | 0.04 |
| MA. 1 (wild) | 0.04 | 0.04 | 0.04 | 0.04 | 0.04 | 0.04 | 0.04 | 0.04 | 0.04 | 0.04 | 0.03 | 0.03 | 0.00 |
| MA. 2 (wild) | 0.82 | 0.81 | 0.81 | 0.81 | 0.81 | 0.81 | 0.82 | 0.82 | 0.81 | 0.82 | 0.58 | 0.58 | 0.00 |
| MA. 3 (wild) | 0.68 | 0.67 | 0.67 | 0.68 | 0.67 | 0.67 | 0.68 | 0.67 | 0.67 | 0.67 | 0.62 | 0.41 | 0.39 |

We found consistent patterns in the efficacy of implementing single-sector alternatives at reducing disease burden in deer. For the agricultural management sector, vaccination and boosting captive deer were the most effective alternatives at reducing disease incidence and persistence in captive settings (Objectives 2 and 3; Table 2). This intervention also had effects on incidence and persistence in sympatric wild deer herds. For public health sector alternatives, we found that increased PPE use during interactions with both captive and wild deer was the most effective alternative but did not have the same magnitude of reduction when compared to vaccination and boosting (Table 2). For wildlife management sector actions, we found that both eliminating baiting or feeding and reducing deer densities were more effective at meeting all measurable attributes for wild deer; however, we found little evidence that those same actions would lead to reductions in disease in captive deer (Table 2).

We then considered the effect of coordinated actions by multiple sectors in line with a One Health approach. Thus, we evaluated the outcome when all sectors implemented their own best performing intervention simultaneously. We found that the combination of vaccination and boosting, use of PPE in captive and wild settings, and the elimination of baiting for wild deer dramatically reduced the median prevalence, incidence proportion, and probability of persistence in all sectors. This effect was larger than the effect sizes that could be achieved by any one sector acting alone (Table 2); this was particularly true for the suburban and intensive captive deer scenario.

## 4. Discussion

The One Health alternative that included actions by all three sectors of governance was the only alternative that eliminated SARS-CoV-2 transmission in captive deer and substantially reduced transmission in wild deer. In general, this finding supports the promise of One Health for improving human, animal, and ecosystem health by jointly considering outcomes in all sectors together. We found that decision analysis was useful in navigating this complex risk and governance problem. By formalizing the problem and providing structure to the decision, this approach helped agencies share information that helped us evaluate SARS-CoV-2 transmission in wild and captive deer and assess the efficacy of alternatives to mitigate it in a way that directly aligns with how agencies view this risk management problem. This study also highlights the potential benefits that decision analysis may provide to agencies in navigating simultaneous decision and governance impediments in disease problems affecting multiple sectors.

While our One Health alternative was drastically more effective, it requires coordination among sectors because of the disjunct authorities to manage different parts of the system. Agricultural agencies have the authority to implement the single most effective alternative: vaccination and boosting. This alternative substantially reduced risk within captive deer and provided benefits to wild deer that may interact along fence lines. While there is currently no vaccine specific to deer and SARS-CoV-2, there are several broad-spectrum vaccines that have proven effective for species that have been inoculated using experimental authority (e.g., Zoetis, Parsippany, NJ, USA). Within the wildlife management sector, reductions in wild deer density and eliminating baiting and feeding reduced disease prevalence and incidence proportion in wild deer. These findings are consistent with other studies that have evaluated the role of density and feeding in intraspecific disease transmission rates^16^. However, lack of compliance with regulations intended to cease baiting and feeding can limit the efficacy of such measures^17^. Finally, we found that public health guidelines and education may be effective if they can increase the proper use of PPE, particularly if PPE is routinely and consistently used during close human-deer encounters.

Our findings provide insight into human-deer spillover and deer-deer transmission in both captive and wild settings under a range of plausible values. While we evaluated conditions that are typical of both deer and human densities in the agro-forested midwestern U.S. (10 deer per km^2^ based on Habib et al. 2011^14^ and Walters et al. 2016^18^; 10-100 humans per km^2^ based on U.S. Census Bureau 2020^19^) there is an opportunity to consider specific combinations of initial conditions and settings to customize results based on measurements from target locations. Based on our settings, we found that human-to-deer spillover was highest in intensive captive settings because of elevated proximity rates estimated by an expert panel^8^. For wild deer, the same panel estimated that suburban settings had 2.7 times higher proximity rates when compared to rural and ranch (captive) deer^9^ (Supplementary Table 3). Despite these differences, Rosenblatt et al. (2023)^8^ found that simulated disease outcomes had low sensitivity across the range of human-to-deer spillover rates, suggesting that infrequent human-to-deer transmission events are sufficient to initiate sustained outbreaks in deer. Deer-to-deer transmission, however, was sensitive to parameter values. The estimated range of R_0_ (0.74 in wild deer and 6.24 in intensive captive deer) was broad and there are many simulations in all settings where R_0_ exceeded 1.0 (this study and Rosenblatt et al. 2023). These findings suggest that while sustained transmission is thought to occur in many wild and captive deer settings^20^, there are likely conditions where deer densities and associated contact rates are insufficient to sustain SARS-CoV-2 outbreaks.

We begin to address simultaneous decision and governance impediments that are common in One Health settings by clearly articulating some of the motivations, mandates, and decisions that each sector is considering. However, to fully understand the tradeoffs in different alternatives and identify practical solutions for SARS-CoV-2 spread in deer, future work would need to evaluate the performance of alternatives across all objectives and potential constraints that are unique to each setting. For example, even though reducing wild deer density was effective, this alternative may not be practical in all areas, especially those with high hunting participation. Creative generation of alternatives, and combination of alternatives that improve the uptake of challenging actions such as reducing deer density, may overcome perceived constraints.

Beyond complex governance, there remain large gaps in our understanding of the ecology and epidemiology of SARS-CoV-2. First, we lack a full understanding of host range. The number of mammalian host species infected with SARS-CoV-2 continues to increase, and with continued evolution, there is potential for host range shifts or expansions. Second, we lack an understanding of how host-pathogen relationships affect intra- and interspecific transmission, spread, and persistence. Although our knowledge of human-to-human spread has improved, many questions remain around spillover, spillback, and long-term pathogen circulation. These uncertainties affect risk mitigation decisions and which governing authorities may be best positioned to act.

One Health actions have proven to be successful at mitigating disease in people, animals, and environments for zoonotic diseases like rabies and Rocky Mountain spotted fever^21^. However, an ongoing challenge is managing the zoonotic spread of SARS-CoV-2 in humans and wildlife because of the complex ways that humans and deer interact in multiple settings and jurisdictions. Here, we applied decision analysis to a One Health problem. We deconstructed the decision problems of multiple sectors into steps that allowed for a partial evaluation of alternatives unique to each sector. Our results can be used in evidence-based decision making and support coordinated, One Health efforts to respond to zoonotic diseases, including SARS-CoV-2.

## Acknowledgments

We thank Margaret McEachran, Jennifer Mullinax, and Michael Tonkovich for their comments and suggestions on previous versions of the manuscript. We thank Casey Barton-Behravesh, Colin Basler, Samantha Gibbs, Colin Gillin, Allen Gosser, Chelsea Gridley-Smith, Paul Johansen, Darlene Konkle, Roxanne Mullaney, Darby Murphy, Susan Rollo, Lisa Shender, Jennifer Siembieda, Jason Sumners, Michael Tonkovich, Nora Wineland. This work was supported by the Coronavirus Aid, Relief, and Economic Security Act (P.L. 116-136). Any use of trade, firm, or product names is for descriptive purposes only and does not imply endorsement by the U.S. Government. This is publication #06 of the Disease Decision Analysis and Research (DDAR) group of the U.S. Geological Survey.

## Disclaimers

The opinions expressed by authors contributing to this journal do not necessarily reflect the opinions of the Centers for Disease Control and Prevention or the institutions with which the authors are affiliated, but do represent the views of the U.S. Geological Survey.

## Data availability statement

All code and generated data used in this study are available in the R software package whitetailedSIRS. Citation: Rosenblatt, E, Rudolph, J.F., Arce, F., Cook, J. D., DiRenzo, G.V., Grant, E.H.C., Runge, M.C., and Mosher, B.A., 2023. whitetailedSIRS: A package to project SARS-CoV-2 outbreak dynamics in white-tailed deer. Version 1.0.0: U.S. Geological Survey software release, https://doi.org/10.5066/P9TZK938

## Notes

https://doi.org/10.5066/P9TZK938

### Competing Interest Statement

The authors have declared no competing interest.

