## Supplementary Materials for "Cross-sector collaboration is more effective than single sector actions at mitigating SARS-CoV-2 in white-tailed deer"

Running title: Cross-sector collaboration reduces SARS-CoV-2 risk in deer

JONATHAN D. COOK^1^, *U. S. Geological Survey, Eastern Ecological Science Center, Laurel, MD, USA*

ELIAS ROSENBLATT, *Rubenstein School of Environment and Natural Resources, University of Vermont, Burlington, VT, USA*

GRAZIELLA DiRENZO, *U. S. Geological Survey, Massachusetts Cooperative Fish and Wildlife Research Unit, University of Massachusetts, Amherst, MA, USA*

EVAN H. CAMPBELL GRANT, *U. S. Geological Survey, Eastern Ecological Science Center, Turner’s Falls, MA, USA*

BRITTANY A. MOSHER, *Rubenstein School of Environment and Natural Resources, University of Vermont, Burlington, VT, USA*

FERNANDO ARCE, *Department of Environmental Conservation, University of Massachusetts, Amherst, MA, USA*

SONJA CHRISTENSEN, *Department of Fisheries and Wildlife, Michigan State University, East Lansing, MI, USA*

RIA R. GHAI, *U.S. Centers for Disease Control and Prevention, Atlanta, GA, USA*

MICHAEL C. RUNGE, *U. S. Geological Survey, Eastern Ecological Science Center, Laurel, MD, USA*

**Supplementary Table 1. Guidance Committee** Names, sector, and affiliations of members of the guidance committee for this study

| **Committee Member** | **Sector** | **Agency** |
| --- | --- | --- |
| Darlene Konkle | Agricultural | Wisconsin Department of Agriculture, Trade, and Consumer Protection |
| Roxanne Mullaney | Agricultural | U.S. Department of Agriculture |
| Susan Rollo | Agricultural | Texas Department of State Health Services |
| Jennifer Siembieda | Agricultural | U.S. Department of Agriculture |
| Nora Wineland | Agricultural | Michigan Department of Agriculture and Rural Development |
| Samantha Gibbs | Wildlife | U.S. Fish and Wildlife Service |
| Colin Gillin | Wildlife | Oregon Department of Fish and Wildlife |
| Allen Gosser | Wildlife | U.S. Department of Agriculture |
| Darby Murphy | Wildlife | U.S. Fish and Wildlife Service |
| Paul Johansen | Wildlife | West Virginia Department of Natural Resources |
| Lisa Shender | Wildlife | National Park Service |
| Jason Sumners | Wildlife | Missouri Department of Conservation |
| Michael Tonkovich | Wildlife | Ohio Department of Natural Resources |
| Casey Barton Behravesh | Public Health | Centers for Disease Control and Prevention |
| Colin Basler | Public Health | Centers for Disease Control and Prevention |
| Ria Ghai | Public Health | Centers for Disease Control and Prevention |
| Chelsea Gridley-Smith | Public Health | National Association of County and City Health Officials |

**Supplementary Table 2.** Complete list of alternative actions that may be effective at mitigating SARS-CoV-2 spread in captive or wild white-tailed deer.

**Alternatives**

1. Manipulate wild populations or habitat conditions.
a. *Local reductions in wild deer populations through agency culling or hunter harvest.*
b. *Prohibit deer feeding and baiting to reduce congregations of wild deer.*
2. Restrictions on research, survey, monitoring or management.
a. *Pause/reduce research permits for work on wild deer or other susceptible wildlife/animal species.*
3. Education and outreach. Intended to communicate the risk that SARS-CoV-2 presents to humans, wildlife, domestic and companion animals. Can be coordinated through veterinarians, public health officials, regulators, hunter’s digest materials, local neighborhood web-apps. Can also be targeted based on human disease trends, wildlife surveillance activities.
a. Encourage proper deer processing activities (field dressing).
b. *Encourage limiting human-deer interactions including participating in conduct that attracts wildlife.*
c. Encourage pet owners to keep animals indoors.
d. *Encourage (and train on the proper) use of enhanced personal protective equipment during interactions with captive and wild deer*
4. Alter/Enhance captive, farming, exhibition, or rehabilitation practices.
a. Depopulation
b. Implement new accreditation program or otherwise enhanced testing
c. *Animal vaccination requirement*
d. *Isolation requirements within facilities or from external environments*
i. *Double fencing*
ii. Enhanced inspections of barriers
e. Limiting interactions between humans and deer (expand focus beyond human-animal welfare to include disease-related enhancements)
f. Limit release of animals
g. Animal food storage and distribution requirements to reduce animal congregations and mixing (biosecurity)
h. *Require enhanced ventilation in enclosed agricultural/zoo/captive settings*
i. Enhanced PPE or vaccination for humans
j. Potential active measures: test and cull, isolate and test, or other restrictions on trade/translocation.
5. Reduce human-wildlife interactions
a. Ban feeding and baiting of wildlife, specifically wild deer, as part of hunting or other wildlife viewing activities.
b. Temporary closures of public trails, parks, other locations.
c. Hazing activities near high-risk areas of human-deer contact, such as wastewater treatment ponds.
d. Regulations on carcass disposal for processors, taxidermists, and other agricultural or hunting activities.

**Supplementary Table 3.** Empirical, expert elicited and fixed parameter values, their sources, and the distributions used to explore parametric uncertainty across the four settings for captive and wild white-tailed deer (reproduced from Rosenblatt et al. 2023).

| Definition (units) | Captive | | Wild | | Source |
| --- | --- | --- | --- | --- | --- |
|  | Outdoor ranch | Intensive facility | Rural | Suburban |  |
| Immune loss rate (day^-1^) | Inverse of log-normal; μ = 4.724, σ = 0.626 | | | | This study, expert elicited |
| Recovery rate (day^-1^) | 1/6 days | | | | Palmer et al. 2021 |
| Proximity rate scaling adjustment (unitless) | 11.35 | NA | 11.35 | 11.35 | Habib et al. 2011 |
| Proximity rate concavity scaling constant (unitless) | 0.34 | NA | 0.34 | 0.34 | Habib et al. 2011 |
| Number of deer per unit area (A_w_) | 1000 | NA | 1000 | 1000 | Habib et al. 2011 |
| Area for intermediate density-dependence (km^2^) | 100 km^2^ | NA | 100 km^2^ | 100 km^2^ | Habib et al. 2011 |
| Adjustment for the presence of an attractant (bait, feed, etc.) | log-normal; μ = 3.473, σ = 0.226 | NA | NA | NA | This study, expert elicited |
| Human-deer proximity rate (events/120 days) | log-normal; μ = 0.572, σ = 0.951 | log-normal; μ = 2.521, σ = 1.132 | log-normal; μ = -1.589, σ = 1.700 | log-normal; μ = 0.572, σ = 0.951 | This study, expert elicited |
| Deer proximity rate in captivity (events/day) | NA | log-normal; μ = 3.470, σ = 0.913 | NA | NA | This study, expert elicited |
| Wild-captive deer proximity rate along fences (events/day, only included for Obejctive 4) | 0.00072 direct contacts/day / σ^DC^ | | | | Vercauteren et al. 2007; Khouri et al. 2022 |
| Quanta SARS-CoV-2 dose-response in deer (1/quanta required for ID63) | log-normal; μ = 0.2775, σ = 0.272 | | | | This study, expert elicited |
| Conversion from SARS-CoV-2 RNA copies to quanta (use human values for deer; quantum/RNA Copy) | 0.0014 quantum/RNA copy | | | | Mikszewski et al. 2021 |
| Concentration of SARS-CoV-2 in human sputum (RNA copies/ml) | μ = 5.6 log10 RNA copies/ml, σ = 1.2 log10 | | | | Buonanno et al. 2020 |
| Concentration of SARS-CoV-2 in deer sputum (RNA copies/ml) | log-normal; μ = 0.216, σ = 0.344; proportional to C_v_ - human | | | | This study, expert elicited |
| Inhalation rate for humans, standing (m^3^/hr) | 0.53 m^3^/hr | | | | Mikszewski et al. 2021 |
| Inhalation rate for deer, breathing (m^3^/hr) | 0.846 m^3^/hr | | | | Ranslow et al. 2014 |
| Droplet volume concentration (speaking; ml/m^3^) | 0.009 ml/m3 | | | | Mikszewski et al. 2021 |
| Volume of shared airspace with 1.5m radius (m^3^) | 7.07 m^3^ | | | | This study, calculated |
| Air exchange rate, recommended rate for hospital rates of 4-6 exchanges per hour (^-hr^) | 4^-hr^ | 1^-hr^ | 4^-hr^ | 4^-hr^ | Gerken et al. 2017; Mathai et al. 2021; Allen and Ibrahim 2021; Bannister et al. 2023 |
| SARS-CoV-2 settling rate (^-hr^) | 0.24^-hr^ | | | | Buonanno et al. 2020 |
| SARS-CoV-2 inactivation rate (^-hr^) | 0.63^-hr^ | | | | Buonanno et al. 2020 |
| Duration of proximity event between human and deer (minutes) | log-normal; μ = 1.788, σ = 1.152 | | log-normal; μ = -0.355, σ = 0.979 | log-normal; μ = 0.432, σ = 0.929 | This study, expert elicited |
| Duration of proximity event between deer (all proximity types; minutes) | log-normal; μ = 1.553, σ = 1.272 | | | | This study, expert elicited |
| Probability of deer making direct contact | logit-normal; μ = -1.457, σ = 0.708 | | | | This study, expert elicited |
| Dose-response function for plaque-forming units (PFU required for ID63) | 410 | | | | Watanabe et al. 2010 |
| Volume of sputum transferred between individuals on contact (μl) | 100 μl | | | | Fixed |
| Concentration of SARS-CoV-2 in deer sputum (RNA copies/ml) | log-normal; μ = 0.216, σ = 0.344; proportional to C_v_ - human | | | | This study, expert elicited |

**Supplementary Methods**

1. **Overview**

To model the spread of SARS-CoV-2 in captive and wild white-tailed deer, we consider direct (aerosolized) and indirect transmission from humans to deer as causing initial deer infections. We then consider direct exposure pathways including: (1) aerosolized SARS-CoV-2 exposure in a shared airspace, and 2) fluid transmission from sputum or other contagious discharges. The time scale is the fall season when most human-deer contact occurs due to hunting activities in the U.S. (September-December).

We started with a simple 2-host system (captive and free-ranging deer), with three compartments Susceptible-Infected-Recovered-Susceptible (SIRS) (Figure 1). Humans were included as a source of infection but were not modeled as a response to disease dynamics in deer.

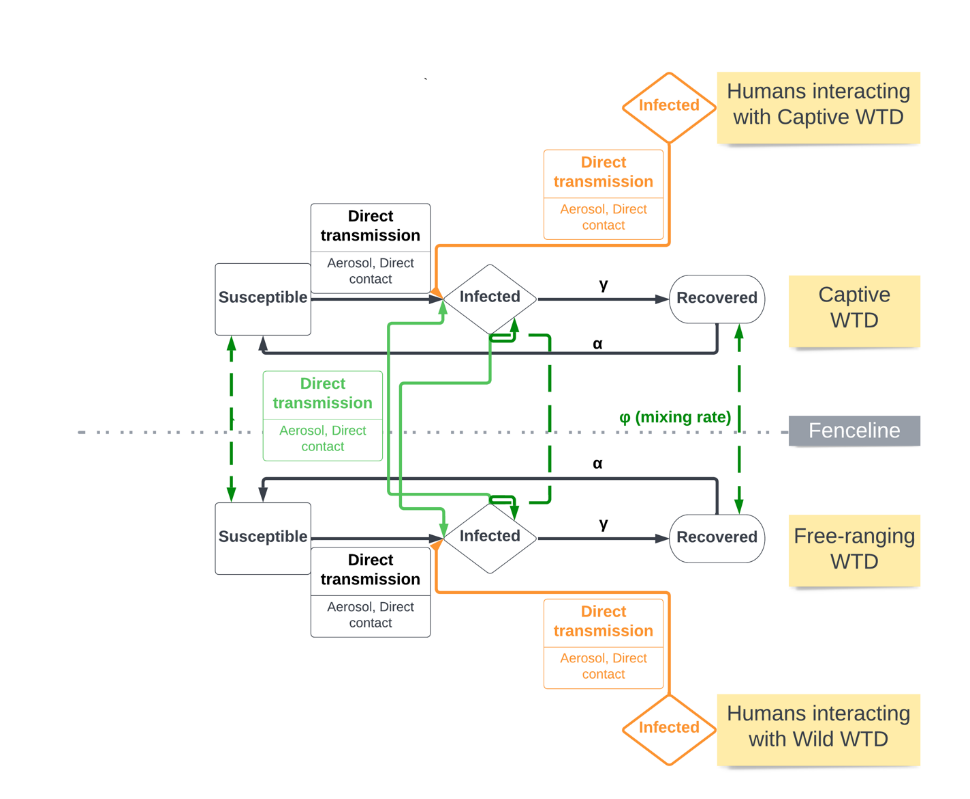

Figure 1: A conceptual diagram of transmission pathways between humans and captive and wild white-tailed deer, facilitated by conspecific interactions (black) within these segments, by fence line interactions or crossings (green), or by interactions with infectious humans (orange) (reproduced from Rosenblatt et al. 2023)

**2.1 Assumptions**

We made the following assumptions for direct transmission pathways:

- transmission rates are additive
- there is homogenous mixing within captive and wild deer populations
- recovery and loss of immunity and mixing within populations do not differ between captive and wild deer
- there is no viral evolution
- there is no disease-induced mortality
- there is no spatial structure
- there is no spillback from deer to humans

**2.2 Model Structure**

We specified an ordinary differential equation (ODE) for the SIRS model (Keeling and Rohani 2007).

Common notation includes:

WW	wild deer to wild deer interaction,

CW	captive deer to wild deer interaction,

WC	wild deer to captive deer interaction,

CC	captive deer to captive deer interaction,

HW	human to wild deer interaction,

HC	human to captive deer interaction,

Aero	aerosolized transmission,

DC	fluid transmission through direct contact.

*2.2.1 Ordinary Differential equation*

We developed three ordinary differential equations to describe disease in wild deer. The daily change in the fraction of susceptible (*S_w_*) deer given by

$$\frac{dS_{w}}{dt}=\alpha R_{w}-S\left( \beta_{WW}^{Aero}I_{W}+\beta_{WW}^{DC}I_{W}+\beta_{CW}^{Aero}I_{C}+\beta_{CW}^{DC}I_{C}+\beta_{HW}^{Aero}I_{H} \right)+\varphi_{CW}S_{C}-\varphi_{WC}S_{w}$$

the daily change in the fraction of the wild population that is infected (*I_w_*) given by

$$\frac{dI_{w}}{dt}=S\left( \beta_{WW}^{Aero}I_{W}+\beta_{WW}^{DC}I_{W}+\beta_{CW}^{Aero}I_{C}+\beta_{CW}^{DC}I_{C}+\beta_{HW}^{Aero}I_{H} \right)+\varphi_{CW}I_{C}-\varphi_{WC}I_{w}- \gamma I_{W}$$

and the daily change in the fraction of wild population that is recovered (*R_w_*) given by

$\frac{dR_{w}}{dt}=\varphi_{CW}R_{C}-\varphi_{WC}R_{W}+\gamma I_{W}-\alpha R_{W}$,

where

α	is the immunity loss rate;

β	is the transmission rate specific to the infectious and susceptible host recipient type (e.g., wild or captive deer) and interactions (i.e., aerosolized or direct contact);

φ	is the intrusion or escape rate (e.g., deer becoming captive or wild);

γ	is the recovery rate from infection.

We also considered captive deer with a similar set of ODEs. The daily change in the fraction of susceptible captive deer (*S_c_*) given by

$$\frac{dS_{C}}{dt}=\alpha R_{C}-S\left( \beta_{CC}^{Aero}I_{C}+\beta_{CC}^{DC}I_{C}+\beta_{CW}^{Aero}I_{W}+\beta_{CW}^{DC}I_{W}+\beta_{HC}^{Aero}I_{H} \right)-\varphi_{CW}S_{C}+\varphi_{WC}S_{w}$$

the daily change in the fraction of the wild population that is infected (*I_C_*) given by

$$\frac{dI_{C}}{dt}=S\left( \beta_{CC}^{Aero}I_{C}+\beta_{CC}^{DC}I_{C}+\beta_{CW}^{Aero}I_{W}+\beta_{CW}^{DC}I_{W}+\beta_{HC}^{Aero}I_{H} \right)-\varphi_{CW}I_{C}+\varphi_{WC}I_{w}- \gamma I_{C}$$

and the daily change in the fraction of wild population that is recovered (*R_C_*) given by

$\frac{dR_{C}}{dt}=-\varphi_{CW}R_{C}+\varphi_{WC}R_{W}+\gamma I_{C}-\alpha R_{C}$.

*2.2.2 Aerosolized Transmission*

Aerosolized transmission rates between a host and recipient ($\beta_{HR}^{Aero}$) is given by

$$\beta_{HR}^{Aero}={\omega_{HR}\times\sigma}^{Aero}$$

where

$\omega_{HR}$	is the proximity rate between host-recipient type (human-wild deer, human-captive deer, wild deer-wild deer, captive deer-captive deer, wild deer-captive deer, captive deer-wild deer);

$\sigma^{Aero}$	is the probability of infection from aerosols.

The proximity rate, $\omega_{HR}$, is based on Habib et al. (2011) and applies to wild deer to wild deer and captive deer to captive deer contact. It is given by

$$\omega_{HR}=\rho_{season}\kappa\left( \frac{{N_{W}}^{\left( 1-q \right)}}{A_{W}} \right)$$

where

$\rho_{season}$	is a seasonal adjustment to density-contact relationship;

$\kappa$	is a scaling constant;

$q$	is a concavity scaling constant;

$N_{W}$	is the total population size;

$A_{W}$	is the area inhabited by the population*.*

All other contact rates (human-wild deer, human-captive deer, wild deer-captive deer, captive deer-wild deer) were included as direct estimates.

The probability of infection, $\sigma^{Aero}$, given proximity is a function of the instantaneous dose received and a Wells-Riley dose response relationship given by

$$\sigma^{Aero}=1- e^{-rQ}$$

where

$r$	is the species-specific probability of infection from 1 quantum of SARS-CoV-2;

$Q$	is the dose received by a single individual.

For reference $r$ = 1 corresponds to 1 quantum causing infection in 63% of susceptible human individuals.

To estimate the dose received by a susceptible individual ($Q$) we modeled 1) the emission of SARS-CoV-2 from an infectious individual (${ER}_{q})$ and 2) the resulting concentration of SARS-CoV-2 in a designated airspace around an infectious individual, considering viral emission and viral loss.

First, an infected individual emits virions at a particular rate (${ER}_{q}$; quanta/hr) as the product of the viral load in its sputum ($C_{v}$; RNA copies/ml), a conversion factor ($C_{i}$; quanta/RNA copy), the inhalation/exhalation rate ($IR$; m^3^/hr), and the exhaled droplet volume concentration ($V_{drop}$; ml droplets/m^3^ exhaled; Mikszewski et al. 2021) given by:

${ER}_{q}=C_{v}\times C_{i}\times IR\times V_{drop}$.

We then use the emission rate to model the instantaneous concentration of virions (C; quanta/m^3^) in a well-mixed airspace ($V_{air}$; m^3^) around an infected individual (${ER}_{q}$; quanta/hr). We assumed that the airspace was 1.0 m^3^. We also account for a loss rate as the sum of air exchange ($AER$; hr^-1^), settling ($s$; hr^-1^), and inactivation ($\lambda$; hr^-1^; modified from Buonanno et al. 2020). Thus, the instantaneous concentration is given by:

$C=\frac{{ER}_{q}}{(AER+s+\lambda)*V_{air}}$*.*

When a susceptible individual enters the contaminated airspace surrounding an infectious individual, the dose ($Q$ ; quanta) is the product of the inhalation rate of the susceptible individual ($IR$; m^3^/hr), the concentration of virions in the fixed volume ($C$; quanta/m^3^), and the duration of contact ($t_{contact}$; hr) given by:

$Q=IR\times C\times t_{contact}$*.*

*2.2.2 Fluid Transmission*

Fluid transmission rates between a host and recipient ($\beta_{HR}^{DC}$) are given by

$$\beta_{HR}^{DC}= \omega_{HR}\times\varepsilon^{DC}\times\sigma^{DC}$$

where

$\varepsilon^{DC}$	is the probability of direct contact;

$\sigma^{DC}$	is the probability of infection from direct contact.

To model fenceline contact between captive and wild deer, we modified eqn. 14 to include a fixed fenceline contact rate, $\omega_{HR}^{fence}$, instead of the contact rate previously described for wild-wild deer and captive-captive deer in eqn. 8.

The probability of infection, $\sigma^{DC}$, given contact was modeled using a Wells-Riley dose response (as in eqn.9) and is given by:

$$\sigma^{DC}=1- e^{-((C_{v}\times V_{sputum})/k)}$$

where

$C_{v}$	is the viral concentration in sputum (in plaque-forming units);

$V_{sputum}$	is the volume of sputum transferred given contact;

$k$	is the dose-response function.
